# Metacognitive ability predicts hippocampal and prefrontal microstructure

**DOI:** 10.1101/046359

**Authors:** Micah Allen, James C. Glen, Daniel Müllensiefen, Dietrich Samuel Schwarzkopf, Martina F. Callaghan, Geraint Rees

## Abstract

The ability to introspectively evaluate our experiences to form accurate metacognitive beliefs, or insight, is an essential component of decision-making. Previous research suggests individuals vary substantially in their level of insight, and that this variation predicts brain volume and function, particularly in the anterior prefrontal cortex (aPFC). However, the neurobiological mechanisms underlying these effects are unclear, as qualitative, macroscopic measures such as brain volume can be related to a variety of microstructural features. Here we used a newly developed, high-resolution (800µm isotropic) multi-parameter mapping technique in 48 healthy individuals to delineate quantitative markers of *in vivo* histological features underlying metacognitive ability. Specifically, we examined how neuroimaging markers of local grey matter myelination, macromolecular and iron content relate to insight. Extending previous volumetric findings, we found that metacognitive ability, as determined by a signal-detection theoretic model, was positively related to the myelo-architectural integrity of aPFC grey matter. Interestingly, perceptual metacognition predicted decreased macromolecule content coupled with increased iron in the hippocampus and precuneus, areas previously implicated in meta-memory rather than meta-perception. Further, the relationship of hippocampal-precuneus and prefrontal microstructure to an auditory memory measure was respectively mediated or suppressed by metacognitive ability, suggesting a dynamic trade-off between participant’s memory and metacognition. These results point towards a novel understanding of the relationship between memory, brain microstructure, and metacognition.

**Significance Statement:** By combining a signal-theoretic model of individual metacognitive ability with state of the art quantitative neuroimaging, our results shed new light on the neurobiological mechanisms underlying introspective insight. Myelination and iron are core determinants of both healthy brain maturation and neurodegeneration; particularly in the hippocampus where iron accumulation is linked to oxidative stress and inflammation. Our results may thus indicate that metacognition depends upon the development and integrity of a memory-related brain network, potentially revealing novel biomarkers of neurodegeneration. These results highlight the power of quantitative mapping to reveal neurobiological correlates of behaviour.

## Introduction

The metacognitive capacity for self-monitoring is at the core of learning and decision-making (Flavell, 1979). As a general capacity, metacognition is thought to enable the flexible monitoring and control of memory, perception and action (Fernandez-Duque et al., 2000). An efficient approach to quantifying this ability lies in the application of signal-detection theory to estimate the sensitivity of self-reported confidence to objective discrimination performance (Fleming and Lau, 2014). Individual differences in metacognitive sensitivity thus quantified are related to the morphological structure, function, and connectivity of the brain (for review, see Fleming and Dolan, 2012). Here we expand on these findings using a newly developed multi-parameter mapping (MPM) and voxel-based quantification (VBQ) technique to better elucidate the neurobiological mechanisms underpinning these effects.

The anterior prefrontal cortex (aPFC) (Fleming et al., 2010, 2012, 2014; Sinanaj et al., 2015) and precuneus (Baird et al., 2013; McCurdy et al., 2013) have been repeatedly related to metacognitive ability. Notably, several studies found a positive relationship between right APFC volume and metacognition (Fleming et al., 2010; McCurdy et al., 2013; Sinanaj et al., 2015). While convergent evidence from anatomical, lesion-based, and functional connectivity studies suggest that the right aPFC is specific to perceptual metacognition, metacognition for memory has instead been related to midline cortical (e.g., mPFC and PCC/precuneus) and hippocampal structures. (Fleming et al., 2012, 2014; Baird et al., 2013; McCurdy et al., 2013). Although these studies suggest that the ability to introspect on perception and memory depends on the development of a neural mechanism involving both domain-specific and general aspects, the underlying neurobiology driving the relationship between neuroanatomy and metacognition remains unclear.

This uncertainty lies partly in the inherent lack of specificity offered by volumetric measures of brain structure, which are fundamentally qualitative in nature. Indeed, voxel-based morphometry (VBM) yields measures in arbitrary units which can be driven by a variety of macroscopic factors such as cortical thickness and variability in cortical folding, owing to a non-specific variety of microstructural features (Ashburner, 2009). It has recently been shown that microstructural properties of brain tissue, such as myelination levels and iron content can lead to the detection of spurious morphological changes (Lorio et al., 2014, 2016). The emerging field of *in vivo* histology aims to combine maps of specific MRI parameters measured via quantitative imaging (qMRI) with biophysical models to provide direct indicators of the microstructural mechanisms driving morphological findings, and ultimately to quantify biologically relevant metrics such as myelination and iron concentrations, oligodendrocyte distributions, and the g-ratio of fibre pathways (Mohammadi et al., 2015; Weiskopf et al., 2015).

In the present study we used qMRI to map a number of key contrast parameters with differential sensitivity to underlying biological metrics, in order to better understand the microstructural correlates of metacognitive ability. To do so, we acquired high-resolution (800µm isotropic) data using the Multi-Parametric Mapping (MPM) qMRI protocol (Weiskopf et al., 2013). Respecting the quantitative nature of these data, we conducted voxel-based quantification (VBQ) analysis (Draganski et al., 2011) in 48 healthy participants to relate these microstructural markers to individual differences in metacognitive sensitivity during an adaptive visual motion discrimination task. Our results confirmed that right aPFC markers of myelo-architecture positively predict metacognitive ability, whereas left hippocampus and precuneus showed effects consistent with both decreased macromolecule and increased iron content. Further clarifying the domain-general role of memory in metacognition, the relationship of hippocampus and precuneus microstructure with auditory memory was mediated by metacognitive ability. These results extend our understanding of the computational neuroanatomy of metacognition and provide novel targets for future clinical research.

## Methods

### Participants

48 healthy participants (29 female) were recruited from University College London and the surrounding community. As age is a strong determinant of micro-architecture and grey matter volume (Callaghan et al., 2014), we restricted our inclusion criteria to 20-40 years, resulting in a mean age of 24 (SD = 5). All participants were right handed, and were mentally and physically healthy with no history of neurological disorders and with normal (or corrected-to normal) vision and hearing. Participants were recruited from a local participant database using broadcast emails. All participants gave informed written consent to all procedures. In accordance with the Declaration of Helsinki, the University College London Research Ethics Committee approved all procedures.

### Study Design

Participants completed the experiment in two sessions, consisting of a 2-hour appointment at the Wellcome Trust Center for Neuroimaging to acquire all imaging data, and a separate 1hour appointment to complete the metacognition task, a brief non-verbal auditory memory measure (Müllensiefen et al., 2014), and other behavioural measures (data not reported here). The neuroimaging session involved 30 minutes of multi-parameter mapping (MPM) while subjects silently viewed a muted nature documentary (to promote stillness). During the behavioural session, participants completed an adaptive psychophysical visual metacognition task (see *Behaviour*, below) lasting 30 minutes.

### Behaviour - Metacognition Global Motion Task

To measure participants’ metacognitive ability, we employed a global dot motion discrimination task comprising a forced-choice motion judgement with retrospective confidence ratings on every trial. As part of another investigation, in which we were investigating noise-induced confidence bias (Spence et al., 2015), we used a dual-staircase approach with two conditions in which either mean direction or standard deviation across dot directions was continuously adapted to stabilize discrimination performance. Thus, to control sensory noise independently of task difficulty, in two randomly interleaved conditions we presented either a stimulus with a fixed 15-degree mean angle of motion from vertical and a variable (adaptive) standard deviation (SD), or a variable (adaptive) mean angle from vertical at a fixed 30 degree SD. In either case, the mean (μ-staircase condition, μ*S*) or standard deviation (σ-staircase condition, σ*S*) of motion was continuously adjusted according to a 2-up-1 down staircase, which converges on 71% performance. On each trial the motion signal was thus constructed using the formula:

DotDirections = Direction (Left vs Right) * MeanOrientation + GaussianNoise*SD

In which the condition-specific staircase determined either mean orientation or SD. Each trial consisted of a 500ms fixation, followed by a 250ms central presentation of the motion stimulus, which was then replaced by a central letter display “L R”. Participants then had 800ms to make their response to indicate whether the mean motion direction was to the left or right of vertical. After this, a confidence rating scale marked by 4 equal vertical lines appeared. Each line was labelled, from left to right “no confidence, low, moderate, high confidence. Participants’ heads were fixed with a chin and forehead rest 72 cm from the screen. Motion stimuli consisted of a central array of 1,100 dots presented over a central fixation dot, within a circular aperture of radius 9.5 degrees visual angle (DVA), with dots advancing 0.02 DVA per frame. To ensure participants attended the global rather than the local motion direction, dot lifetimes were randomized and limited to a maximum of 93% stimulus duration.

**Figure 1.**
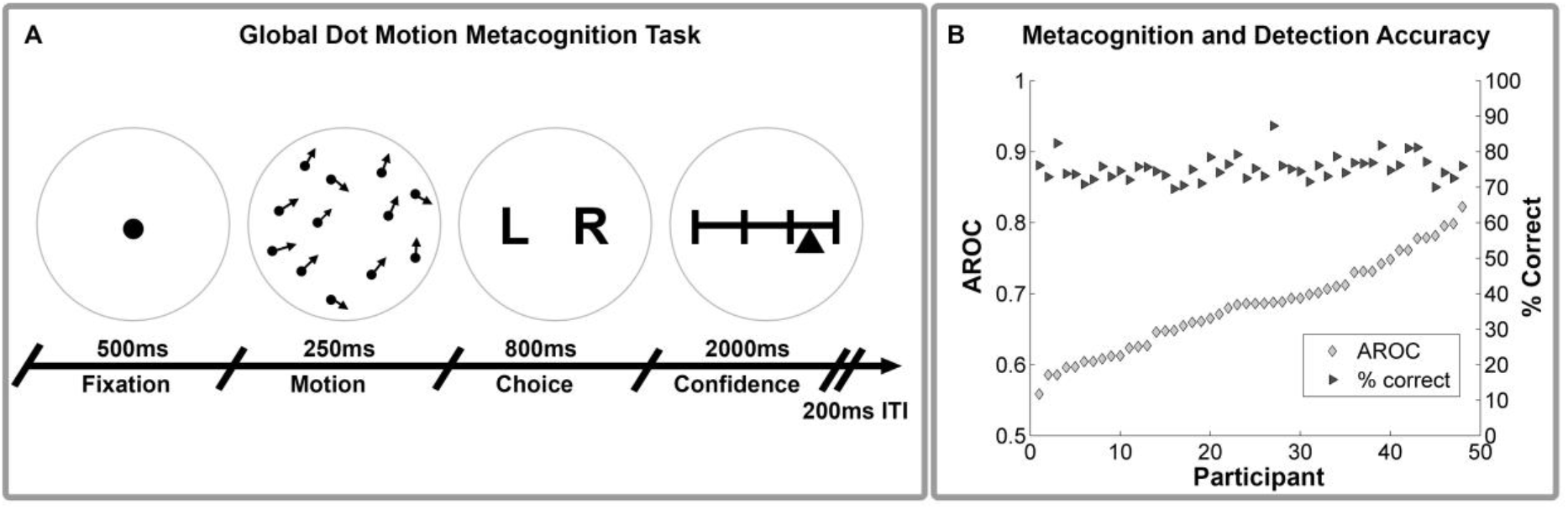
**Behavioral Paradigm and Metacognitive Accuracy**. Schematic of global dot-motion metacognition task (A) and plot of metacognitive vs motion detection accuracy (B). Participants were required to judge the global or average motion of a brief dot display, and then rate their confidence in this judgement from 0 (guessing) to 100 (certain). Performance was held constant using an adaptive threshold adjusting either signal mean or variance on each trial (see *Methods* for more details). Right hand plot demonstrates substantial individual differences in metacognitive accuracy, estimated as the type-II area under the curve (AROC), independently of motion discrimination performance. Inter-individual differences in AROC were then used in a multiple regression analysis to explain variation in microstructural brain features (see *VBQ Analysis*).

Participants were instructed that the goal of the task was to measure their perceptual and metacognitive ability. Metacognitive ability was defined as a participant’s insight into the correctness of their motion judgements, i.e. how well their confidence reports reflected their discrimination accuracy. Participants completed a short practice block of 56 trials, in which they performed the motion discrimination without confidence ratings, with choice accuracy feedback provided by changing the colour of the fixation to green or red. All participants achieved better than 70% accuracy and indicated full understanding of the task before continuing. Participants completed 320 trials divided evenly between the two staircase conditions. Trials were divided into 10 blocks each with 40 trials, randomly interleaved across conditions within each block. 14 participants did not complete the last two blocks of the task due to a technical error, however all participants had at least 100 trials per condition (Fleming and Lau, 2014).

### Non-Verbal Auditory Measure

As previous studies have found both commonalities and differences between metacognition for perception and memory (Baird et al., 2013; McCurdy et al., 2013), we collected a brief measure of participant’s non-verbal auditory memory using a common same-different comparison paradigm (Müllensiefen et al., 2014). This involved a short AB task in which participants listened to a sequence of 10-17 notes, and then decided if a subsequent sequence at a different absolute pitch level had the same or a different structure. The second note sequence was always transposed either by a fifth or by a semitone. Participants were required to indicate whether the two note sequences had an identical pitch interval structure or not. The test score was then calculated as the accuracy of the same-different judgement for each participant across all 13 trials.

### Quantitative Multi-Parameter Mapping (MPM)

Recent technical developments have enabled *in vivo* mapping of neuroimaging markers of biologically relevant quantities to be performed with high resolution and whole brain coverage (Deoni et al., 2005; Helms et al., 2008a, 2008b, 2009). We used the Multi-Parameter Mapping (MPM) protocol (Weiskopf et al., 2013) to obtain maps of the percent saturation due to magnetization transfer (MT), longitudinal relaxation rate (R_1_), and effective transverse relaxation rate (R_2_*).

### Data Acquisition

All imaging data were collected on a 3T whole body MR system (Magnetom TIM Trio, Siemens Healthcare, Erlangen, Germany) using the body coil for radio-frequency (RF) transmission and a standard 32-channel RF head coil for reception. A whole-brain quantitative MPM protocol consisting of 3 spoiled multi-echo 3D fast low angle shot (FLASH) acquisitions with 800 µm isotropic resolution and 2 additional calibration sequences to correct for inhomogeneities in the RF transmit field (Lutti et al., 2010, 2012; Callaghan et al., 2015b).

The FLASH acquisitions had predominantly proton density (PD), T1 or MT weighting. The flip angle was 6^0^ for the PD- and MT-weighted volumes and 21^0^ for the T1 weighted acquisition. MT-weighting was achieved through the application of a Gaussian RF pulse 2 kHz off resonance with 4ms duration and a nominal flip angle of 220^0^. The field of view was 256mm head-foot, 224mm anterior-posterior (AP), and 179mm right-left (RL). Gradient echoes were acquired with alternating readout gradient polarity at eight equidistant echo times ranging from 2.34 to 18.44ms in steps of 2.30ms using a readout bandwidth of 488Hz/pixel. Only six echoes were acquired for the MT-weighted acquisition in order to maintain a repetition time (TR) of 25ms for all FLASH volumes. To accelerate the data acquisition, partially parallel imaging using the GRAPPA algorithm was employed with a speed-up factor of 2 in each phase-encoded direction (AP and RL) with forty integrated reference lines.

To maximise the accuracy of the measurements, inhomogeneity in the transmit field was mapped using the 2D STEAM approach described in Lutti *et al*. 2010, including correcting for geometric distortions of the EPI data due to B0 field inhomogeneity. Total acquisition time for all MRI scans was less than 30 mins.

### Parameter Map Estimation and Voxel-Based Quantification (VBQ)

All images were processed using SPM12 (version 12.2, Wellcome Trust Centre for Neuroimaging, http://www.fil.ion.ucl.ac.uk/spm/) and bespoke tools implemented in the voxel-based quantification (VBQ) toolbox version 2e (Draganski et al., 2011; Weiskopf et al., 2015), implemented in MATLAB (Mathworks Inc, version R2014a).

To create the quantitative maps, all weighted volumes were co-registered to address inter-scan motion. Maps of R_2_* were estimated from the gradient echoes of all contrasts using the ordinary least squares ESTATICS approach (Weiskopf et al., 2014). The image data for each acquired weighting (PDw, T1w, MTw) were then averaged over the first six echoes to increase the signal-to-noise ratio (SNR) (Helms et al., 2009). The three resulting volumes were used to calculate MT and R_1_ as described in (Helms et al., 2008a, 2008b) including corrections for transmit field inhomogeneity and imperfect spoiling (Preibisch and Deichmann, 2009a, 2009b; Callaghan et al., 2015c). The MT map depicts the percentage loss of signal (MT saturation) that results from the application of the off-resonance MT pre-pulse and the dynamics of the magnetization transfer (Helms et al., 2008b).

A Gaussian mixture model implemented within the unified segmentation approach was used to classify MT maps into grey matter (GM), white matter (WM) and cerebrospinal fluid (CSF) (Ashburner and Friston, 2005). Diffeomorphic image registration (DARTEL) was used to spatially normalise individual grey and white matter tissue classes generated from the structural MT maps to a group mean template image (Ashburner, 2007). The resulting DARTEL template and participant-specific deformation fields were used to normalise the MT, R_1_ and R_2_* maps of each participant to standard MNI space. We based our normalization of the quantitative map on the MT maps because of their greatly improved contrast in subcortical structures, e.g. basal ganglia, and similar WM/GM contrast in the cortex to T1-weighted images (Helms et al., 2008a). A 4 mm full-width at half-maximum (FWHM) Gaussian smoothing kernel was applied to the R_1_, MT, and R_2_* during normalisation using the VBQ approach, which aims to minimise partial volume effects and optimally preserve the quantitative values (Draganski et al., 2011). This tissue-specific approach to smoothing generates grey and white matter segments for each map. The grey matter segments were used in all subsequent analyses. For results visualization, an average MT map in standard MNI space was created from all participants.

## Analysis

### Behavioural Analysis – Detection Performance and Metacognitive Ability

All behavioural data were pre-processed using MATLAB (The Mathworks Inc, Natick, MA, USA). Following previous investigations, we discarded the first block of trials to allow for staircase stabilization. Any trial with reaction times (RT) below 100ms or more extreme than 3 SD of mean RT was rejected from analysis. To quantify metacognitive ability, we estimated the type-II area under the receiver-operating curve (AROC) (Fleming et al., 2010, 2012; Fleming and Lau, 2014) separately for each staircase condition. This method has been described extensively elsewhere (Fleming et al., 2010; Fleming and Lau, 2014), but briefly it involves constructing a receiver-operating curve describing the p(high confidence | correct response) vs the p(low confidence | incorrect response). Under equal performance the AROC thus describes the sensitivity of a participant’s confidence ratings relative to their actual performance. For AROC estimation, confidence ratings were binned into 4 equally sized quartiles (a necessary step for fitting the signal-theoretic model) using bespoke MATLAB code. As a general index of metacognitive ability, we then calculated average AROC, as well as average confidence, mean accuracy (% correct responses), detection sensitivity (*d’*), choice bias (*c*), and reaction time (*RT*). We also calculated median signal mean, median signal variance, and accuracy within each condition to characterize our thresholding procedure.

### VBQ Analysis – Metacognitive Ability

We initially focused on replicating and extending previous volumetric findings relating metacognitive ability to neuroanatomy (Fleming et al., 2010, 2014; McCurdy et al., 2013, 2013; Sinanaj et al., 2015). To do so we conducted volume of interest (VOI) multiple regression analyses using 5mm-radius spherical VOIs centred on the peak coordinates reported by Fleming (2010) and McCurdy (2013). This involved the creation of VOIs in the left (−20, 53, 12), (−12, 54, 16) and right (24, 65, 18), (33, 50, 9), (32, 50, 7) APFC, right dorsolateral PFC (36, 39, 21), and precuneus (6, −57, 18), (8, −64, 24). All VOIs were generated using the WFU Pickatlas Version 3.0.5 (Maldjian et al., 2003), and combined into a single mask. The resulting mask is available for download at Neurovault.org (http://neurovault.org/collections/1260/). We then modelled inter-subject variation in the MT, R_2_*, and R_1_ grey matter maps in separate random-effects multiple regression analyses, modelling average AROC as our key dependent variable. Importantly, we followed previous investigations and controlled all analyses for average discrimination sensitivity (d-prime), confidence, response bias, the variance-induced confidence bias, and the difference in mean signal between the two staircases. To estimate variance-induced confidence bias, we fit multiple regression models within each subject, modelling trial-wise mean, variance, accuracy, and RT as predictors of confidence. This provided beta-weights for each participant indicating the degree to which their confidence report reflected variance-specific bias independently of the other modelled factors, which were then included in our VBQ multiple regression.

Following standard VBM procedure, we also included age, gender, and total intracranial volume as nuisance covariates. We then conducted small-volume corrected analyses of the positive and negative main effect of metacognitive ability (AROC) within our *a priori* mask, correcting for multiple comparisons using Gaussian Random Field Theory, FWE-peak corrected alpha = 0.05. Further, we analysed the whole-brain maps of the same contrasts, using a non-stationarity corrected FWE-cluster p-value with a p < 0.001 inclusion threshold (Ridgway et al., 2008; Hupé, 2015). All anatomical labels and % activations were determined using the SPM Anatomy Toolbox (Eickhoff et al., 2005).

### Mediation Analysis of Auditory Memory and Metacognition

To explore the relationship of metacognition, memory, and brain microstructure we conducted a single-level mediation analysis with the auditory memory score (*X*) predicting brain microstructure (*Y*), mediated by metacognitive ability (*M*) (Wager et al., 2009; Woo et al., 2015). This enabled us to infer whether the relationship of, e.g., hippocampal microstructure and memory depended upon metacognition. For this analysis, R_2_* and MT values were extracted from each subject within the peak coordinate identified by our multiple-regression analysis, for the APFC (MT only), precuneus, and left hippocampus (see *VBQ Results*, below). As we observed a similar pattern of correlation for AROC in both hippocampus and precuneus, we first assessed the degree of correlation between each region for R_2_* and MT, respectively. Indeed, R_2_* and MT were significantly correlated between both regions (R_2_* between region R = 0.37, p = 0.009; MT between region R = 0.46, p = 0.002). We thus averaged the values of each parameter across regions to create a general hippocampal-precuneus (HP) index for both measures. We then fit single-level mediation analyses using the Multilevel Mediation and Moderation Toolbox for MATLAB (https://github.com/canlab/MediationToolbox) to HP R_2_*, HP MT, and APFC MT while controlling for the same covariates as in our *VBQ analysis*. Group-level significance for all direct (*a*,*b*,*c*,*c’*) and mediation parameters (*ab*) was assessed using a bootstrapping procedure with 10,000 bootstrap samples, alpha = 0.05.

## Results

### Behavioural Results

To check staircase stability, we first performed two-way repeated measures ANOVA (factor A: block, levels 1-7; Factor B: staircase condition, μ*S vs* σ*S*) on accuracy scores after removing the first block. As several subjects did not complete the last two blocks, we first re-binned trials into 8 equal size bins of 20% total trial length, before analysing block stability. This analysis revealed a significant main effect of variance on accuracy (*F*(1, 47) = 15.15, *p* < 0.001), but no main effect of block (*ps* > .33) or block by condition interaction (*p* > 0.11), indicating that although average performance was slightly higher in the σ*S* condition (Mean Accuracy μ*S* = 73.4%, Mean Accuracy σ*S* = 76.6%), this difference did not change over time, indicating stable performance within each staircase. As a further check we repeated this analysis separately within each condition; in both cases the block main effect was not significant (all *ps* > 0.13). All participants thus achieved stable performance, with an average accuracy of 75.3% (SD = 3%) across two conditions. Metacognitive ability was comparable to previous studies using the AROC (mean AROC = 0.68, SD = 0.06) and did not differ between conditions, *t* (47) = −0.67, p = 0.51. Table 1 presents descriptive statistics for discrimination and metacognition performance.

**Table 1.** **Behavioural Summary**

| Variable | $\mu S$ (SD) | $\sigma S$ (SD) | $\mu (\mu S, \sigma S)$ (SD) |
| --- | --- | --- | --- |
| d' | 1.40 (0.32) | 1.60 (0.41) | -0.20 (0.43) |
| RT (ms) | 404.3 (66) | 414.0 (69) | 409.1 (66) |
| Accuracy % | 73.4 (3.6) | 76.8 (4.9) | 75.3% (3) |
| Confidence (1-4) | 2.75 (0.26) | 2.39 (0.27) | 2.56 (0.14) |
| Confidence (1-100) | 73.4 (13.8) | 62.6 (15.2) | 62.64 (15.23) |
| AROC | 0.68 (0.07) | 0.69 (0.08) | 0.68 (0.06) |
| Median $\mu$ or $\sigma$ | 6.46 (2.96) | 53.34 (13.27) | na |

Note: $\mu S$ = mean staircase, $\sigma S$ = variance staircase, SD = standard deviation.

### Neuroimaging Results

#### VOI analysis - extension of previous APFC and precuneus Findings

Our VOI analysis revealed significant correlations within the right precuneus and aPFC. aPFC showed overlapping, significant positive correlations of AROC with both R_1_ (peak voxel MNI_xyz_ = [37 41 22]) and MT (peak voxel MNI_xyz_ = [37 42 22]) maps. As both maps are sensitive to myelination to a varying degree, this suggests previous volumetric findings in the aPFC are related to the myelo-architecture of the cortical grey matter. In contrast, in the precuneus we found that AROC was negatively related to MT (peak voxel MNI_xyz_ = [10 −58 8]) and positively related to R_2_* (peak voxel MNI_xyz_ = [9 −64 24]). This result likely indicates a complex interaction between iron-regulating macromolecules such as ferritin, microglia, or amybetaloids and local iron concentration (Galazka-Friedman et al., 2009). Interestingly, it may explain why some previous studies failed to find correlates of perceptual metacognition in the precuneus (Fleming et al., 2010; Sinanaj et al., 2015), as this pattern of microstructural changes may counteract resulting in minimal contrast on T1-weighted images (Callaghan et al., 2015a) and therefore no net difference in a VBM analysis using such images. See table 2 and Figure 3 for summary of these results.

**Figure 2.**
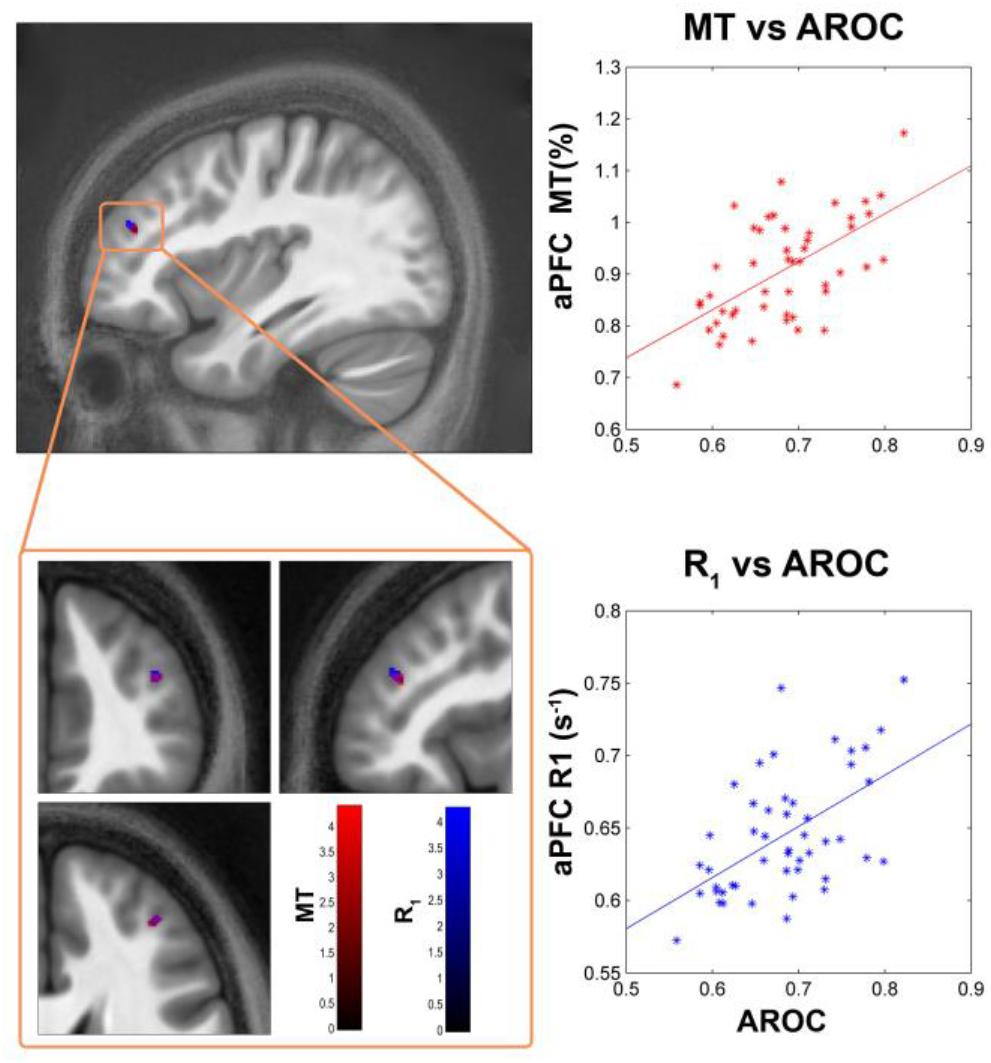
**aPFC VBQ findings**. Figure shows correlation of metacognitive ability (AROC) and anterior prefrontal (aPFC) microstructural measures of white-matter concentration (R_1_ and MT) across 48 participants. Right side, scatter plots showing peak voxel vs AROC, with least-squares line for illustration purposes. Bottom left, zoomed in view shows overlap of AROC correlation in both MT and R_1_ maps. Colour bars indicate t-values, blobs displayed on average MT map from our 48 participants. Volume of interest analysis, FWE-peak corrected p < 0.05 within mask generated from previously reported coordinates (Fleming et al., 2010; McCurdy et al., 2013). See *VBQ Analysis* for more details.

**Table 2.** **Summary of Brain Results**

VOI Results
| Map | Region | k | p<sub>FWE</sub> peak | p<sub>U</sub> peak | T | Z | x | y | z |
| --- | --- | --- | --- | --- | --- | --- | --- | --- | --- |
| MT + | R APFC | 38 | 0.035 | < 0.001 | 4.38 | 3.92 | 37 | 41 | 22 |
| MT - | R Precuneus | 25 | 0.024 | < 0.001 | 4.54 | 4.03 | 10 | -58 | 8 |
| R<sub>2</sub>* + | R Precuneus | 51 | 0.040 | < 0.001 | 4.20 | 3.78 | 9 | -64 | 24 |
| R<sub>1</sub> + | R APFC | 51 | 0.043 | < 0.001 | 4.25 | 3.82 | 37 | 42 | 22 |

Whole Brain Results
| Map | Region | k | p<sub>FWE</sub> cluster | p<sub>U</sub> cluster | T | Z | x | y | z |
| --- | --- | --- | --- | --- | --- | --- | --- | --- | --- |
| MT - | L Hippo. | 850 | 0.030 | < 0.001 | 5.48 | 4.68 | -31 | -25 | -14 |
| R<sub>2</sub>* - | L & R V1 | 598 | 0.016 | < 0.001 | 4.89 | 4.28 | 1 | -69 | 11 |
| R<sub>2</sub>* + | L MTG | 568 | 0.011 | < 0.001 | 5.72 | 4.83 | -51 | -48 | 2 |
| R<sub>2</sub>* + | R Subiculum<sup>†</sup> | 417 | 0.044 | < 0.001 | 6.17 | 5.10 | 14 | -38 | -5 |
| R<sub>2</sub>* + | L Hippo.<sup>‡</sup> | 5518 | < 0.001 | < 0.001 | 4.94 | 4.31 | -26 | -36 | -13 |

Note: Hippo = hippocampus, MTG = middle temporal gyrus, APFC = rostralateral prefrontal cortex. <sup>†</sup> indicates peak-corrected result and p-values. <sup>‡</sup> indicates result from exploratory $p < 0.01$ inclusion threshold. $k$ = cluster extent in voxels, T = t-value, Z = z-value, x,y,z = MNI peak coordinates. pFWE cluster = family-wise corrected cluster p-value, pU cluster = uncorrected cluster p-value, pFWE peak = family-wise corrected peak p-value, pU cluster = uncorrected peak p-value. + - indicates positive or negative t-contrast. See *VBQ analysis* and *VBQ Results* for more details.

#### Whole-brain AROC Analysis

Our whole brain analysis revealed a striking relationship between AROC and left hippocampal MT. Here, higher AROC related to reduced MT in the left posterior-hippocampus (peak voxel MNI_xyz_ = [−31 −25 −14]). Inspection of this result in the SPM anatomy toolbox revealed that the majority (51.6%) was in the dentate gyrus (33.4% ‘activated’), with another 29.3% in CA1 (14.2% activated), and to a lesser extent in the subiculum (6.6%, 1.8% activated) and CA3 (5.6%, 14.9% activated). Additionally, we found that iron levels as indexed by R_2_* negatively predicted AROC in the visual cortex, primarily in V1 (peak voxel MNI_xyz_ = [1 −69 11]), and positively predicted AROC in the left middle-temporal gyrus (peak voxel MNI_xyz_ = [−51 −48 2]). A positive iron effect which was just above our cluster-level threshold (pFWE cluster = 0.052), but which was FWE-peak significant (pFWE peak = 0.044) was also found in the right subiculum (peak voxel MNI_xyz_ = [14 −38 −5]). Finally, as our VOI analysis indicated an inverse relationship between precuneus MT and R_2_*, we were interested to see if a similar pattern could be found in the left hippocampus at a reduced (i.e., more exploratory) threshold. We thus lowered our uncorrected inclusion threshold to p < 0.01, FWE-cluster corrected to p < 0.05, and found that left hippocampal R_2_* also positively predicted AROC (peak voxel MNIxyz = [−26 −36 −13], pFWE < 0.001). These results are particularly interesting in light of the fact that they have not been previously reported in the context of perceptual metacognition, which may be due to the fact that decreased macromolecular content coupled with increased iron content may result in no net volume change being identified, highlighting the increased sensitivity and specificity of our method. All raw t-maps and FWE-thresholded maps have been made available to view and freely downloaded at Neurovault (http://neurovault.org/collections/1260/).

**Figure 3.**
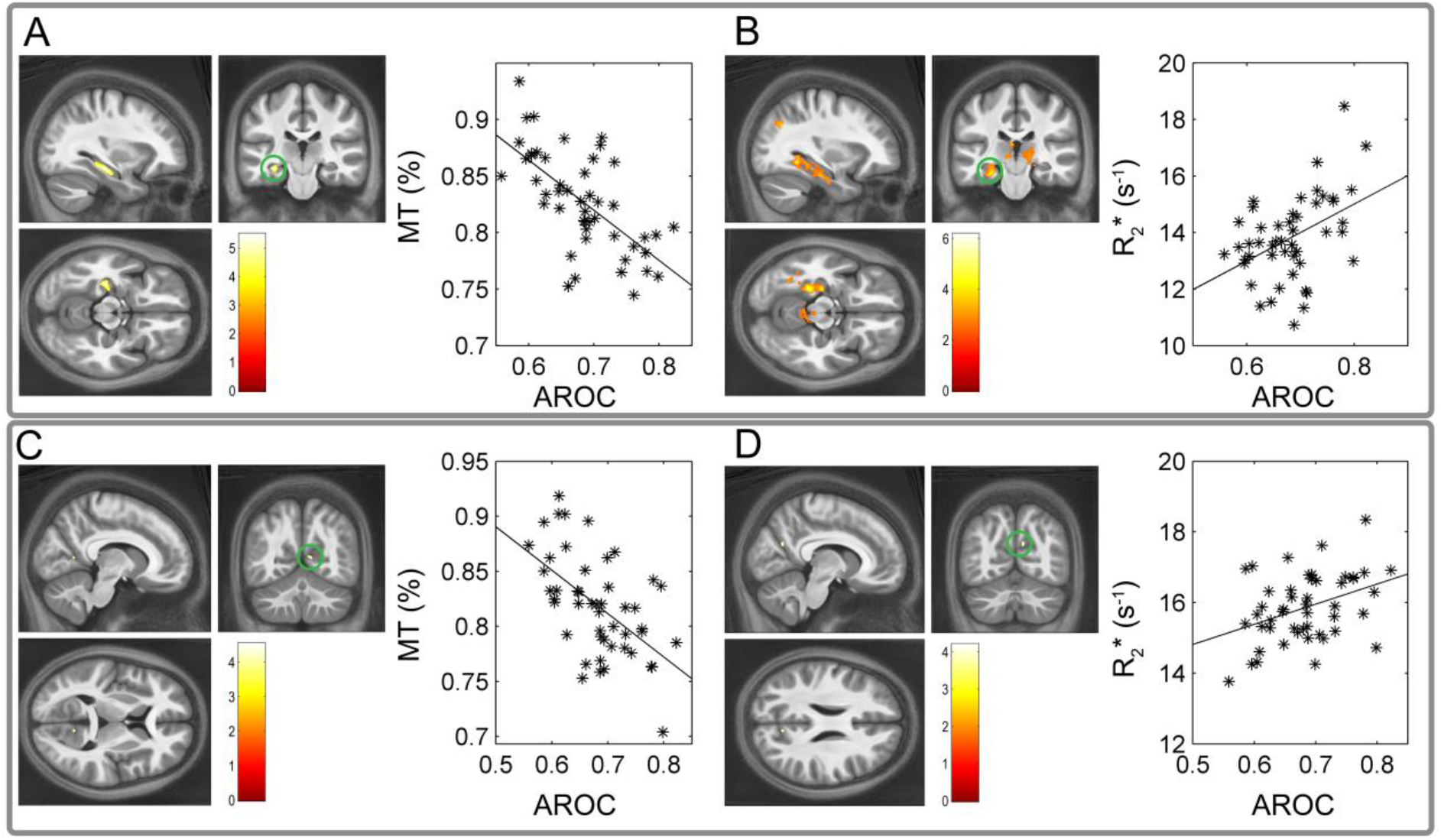
Hippocampus and Precuneus VBQ findings. Voxel-based quantification results in the left hippocampus and right precuneus. Hippocampus MT negatively relates to metacognitive ability (AROC) (top left, A), whereas iron levels in the same region positively predict metacognition (top right, B). Interestingly, the same pattern is seen for right hippocampus. Blobs depict results of multiple regression analyses vs each map type, while controlling for age, gender, ICV, and a variety of performance-related variables (see *Methods* for more information). Results shown on average MT map in MNI space. Scatterplots are for illustration purposes only and depict the peak voxel from each SPM versus raw AROC. Top panel results are from FWE-cluster corrected whole brain analysis, pFWE < 0.05; bottom panel results are from VOI analysis across multiple regions drawn from previous volumetric metacognition studies.

#### Mediation Results - Auditory Memory and Metacognition

This analysis revealed a significant relationship between memory scores and AROC (mean path = 0.39, SD = 0.17, p < 0.05). Across all three brain measures, AROC significantly mediated the relationship of brain microstructure and memory ability – interestingly for MT this relationship was positive for the aPFC, with higher metacognition mediating the relationship of memory and aPFC (mean path = 0.24, SD = 0.15, p < 0.05); instead suppression (negative mediation) was observed for hippocampal-precuneus (mean path = −0.34, SD = 0.16, p < 0.05). Finally, for HP R_2_* AROC also showed a positive mediation effect (mean path = 0.27, SD = 0.16, p < 0.05). These results suggest that metacognition differentially changes the interrelation of prefrontal and hippocampal microstructure to memory ability, perhaps indicating that individuals who are better at metacognition have a more ‘metacognitive’ memory network and conversely, a more memory-related metacognition system. See figure 4 for an overview of these results.

**Figure 4.**
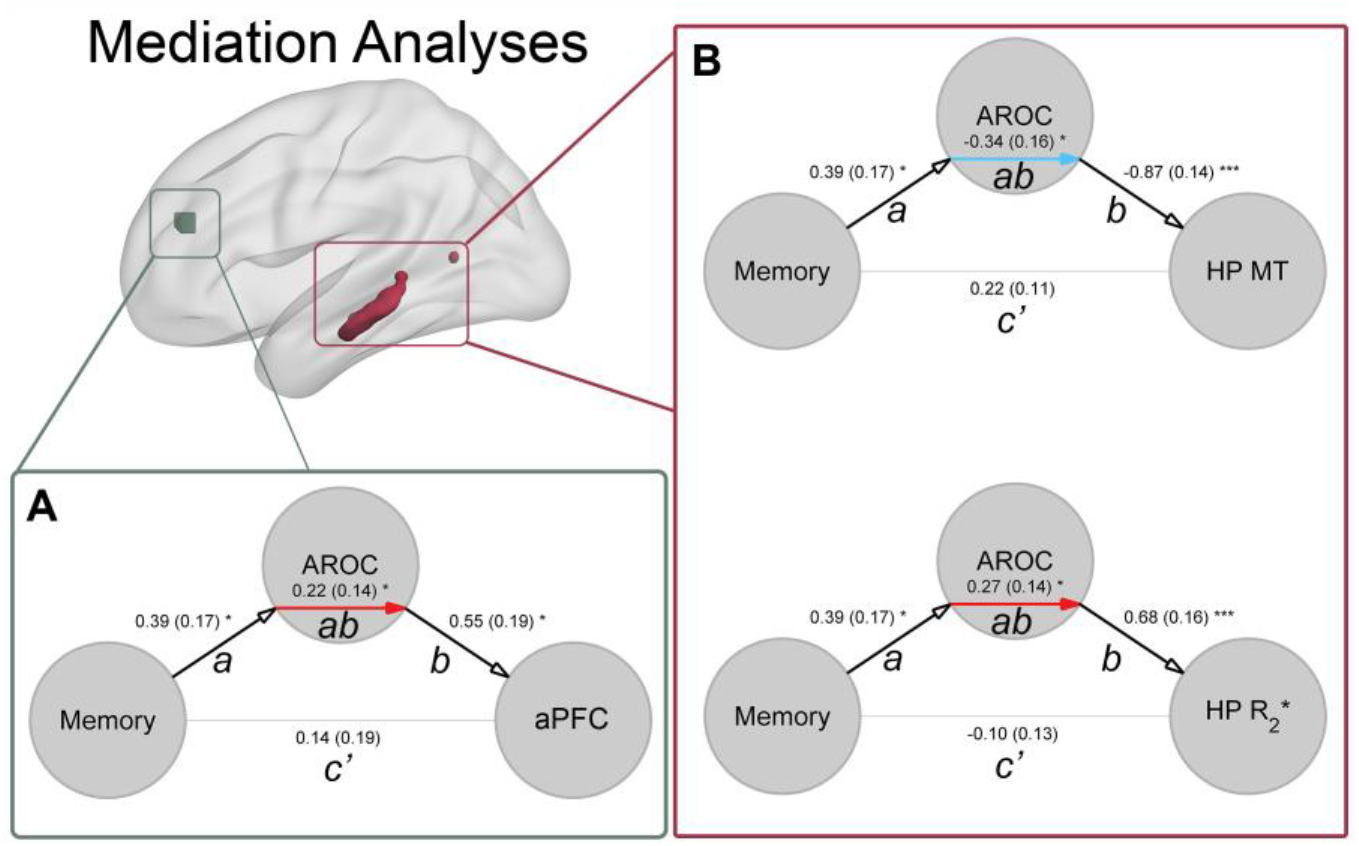
**Memory and Metacognition Mediation Analysis Results**. **A)** Metacognitive ability (AROC) significantly mediates (red arrow) the relationship of non-verbal auditory memory and APFC myelo-architecture (as measured by Magnetization Transfer Ratio, MT). In contrast, **B)** the relationship of memory and hippocampal-precuneus (HP) myelo-architecture is supressed by metacognitive ability (blue arrow), but enhanced for HP iron (as measured by R_2_*). These results suggest individual differences in memory and metacognition are related by a dynamic interaction between the two networks, with the memory-related network increasingly related to metacognitive circuits and vice versa for the prefrontal cortex. Statistical significance for each parameter of the path model (*a*, *b*, *ab*, and *c’*; * < 0.05, *** < 0.001) determined via bootstrapping procedure, see *Methods* for more details.

## Discussion

Our findings demonstrate that individual differences in metacognitive ability are related to underlying microstructural features of prefrontal and hippocampal neuroanatomy. Previous studies investigating individual metacognitive ability indicated that the function, connectivity, and volume of anterior prefrontal cortex (aPFC) underlie introspective insight. Here we build on these findings using a novel quantitative magnetic resonance technique to show that prefrontal correlates of metacognition are related to markers of grey-matter myelo-architecture, as indexed by the overlapping effect in both MT and R_1_ maps. In contrast, we found that differences consistent with decreased hippocampus and precuneus macromolecular content, coupled with increased iron content predicted metacognitive ability. Interestingly, we also found that metacognition mediated the relationship of memory ability and microstructure in the hippocampus and prefrontal cortex, suggesting a domain-general mechanism linking memory and metacognition. These results suggest that unique neurobiological mechanisms underlie the development and aetiology of metacognition for memory (i.e., meta-memory) and perception (meta-perception).

Previous investigations of metacognitive ability and individual differences in brain anatomy suggest that introspection for memory and perception depend upon both common and unique neural substrates. As strong evidence for their dissociation, Fleming et al (2014) recently demonstrated that medial-prefrontal (MPFC) or aPFC lesions selectively disrupt meta-memory or perception, respectively. In another study by McCurdy et al (2013), although metacognition for perception and memory were found to correlate, the two processes independently related to the volume of aPFC and precuneus. Interestingly, in an interaction analysis these authors found that while precuneus volume also predicted meta-perceptual ability, aPFC did not predict meta-memory. In contrast, Baird and colleagues (2013) found no behavioural correlation of meta-memory and perception, but instead found that the former predicted functional connectivity from the medial prefrontal cortex (mPFC) to the precuneus and inferior parietal cortex, while the latter was predicted by functional connectivity seeded from the aPFC to the cingulate, putamen, and thalamus. However, Baird et al. also examined the differential connectivity strength of aPFC vs mPFC and found that better meta-memory was associated with stronger connectivity between the aPFC and the hippocampus, precuneus, and other memory-related areas, a finding which may possibly explain our mediation results. Thus while these studies suggest a degree of independence between meta-memory and meta-perceptual systems, they also suggest that interactions between memory (precuneus, hippocampus) and metacognition-related (aPFC) brain areas underlie general metacognitive ability.

Complementing these results, we found that perceptual metacognitive ability was positively related to the myelo-architectural integrity of the aPFC, as evidenced by coincident AROC-related differences in R_1_ and MT maps in this region. In contrast, in the precuneus and left hippocampus we found that decreased MT and increased R_2_* were predictive of inter-individual differences in perceptual metacognition. Furthermore, the relationship of microstructure in each of these areas to memory was mediated by metacognitive ability, supporting the hypothesis of an interaction between memory and perceptual awareness. Our results are thus consistent with a general linkage between memory, metacognition, and related neural networks.

Although each of the multi-parameter maps have enhanced specificity over conventional weighted imaging and each exhibit sensitivity to particular microstructural tissue properties, this relationship is not unique. Histologically, differences in MT measures strongly correlate with myelin content (e.g., Mottershead et al., 2003; Schmierer et al., 2007; Turati et al., 2015). While myelin is also a significant determinant of R_1_ (Koenig, 1991; Mottershead et al., 2003; Gouw et al., 2008), other features of the myelo-architecture such as the axonal diameter, perhaps coupled to the exposed myelin surface, may be a greater determinant, at least in white matter (Harkins et al., 2016). Nonetheless, other factors such as iron content (Gelman et al., 2001; Rooney et al., 2007; Callaghan et al., 2015a) and cellular architecture (Mottershead et al., 2003; Gouw et al., 2008) also play a role. R_2_* is highly correlated with iron content (Langkammer et al., 2010) but can also be influenced by fibre orientation and architecture (Wharton and Bowtell, 2012). Considering this array of factors, we benefit from having a multi-modal view to aid in interpreting our findings. Thus, where we identified a decreased MT coupled with an increased R_2_* in the absence of an R_1_ effect suggests a coincident decrease in myelin and increase in iron content, potentially driven by changes in underlying microglial and astroglial support structures. In contrast, the co-localisation of increases in both MT and R_1_ suggests an increase in myelination independent of changes in iron or other paramagnetic content that would also be expected to impact R_2_*.

This pattern of results may suggest a unique neurobiological mechanism for the interaction of stress, memory, and metacognition. One recent study by Reyes et al (2015) tested individual response to stress induction, and then at a later date participant’s perceptual metacognitive ability. These authors found that high-stress responders showed an altered overall level of confidence and reduced metacognitive ability. This is particularly interesting in light of our findings in the left hippocampus. The hippocampus is highly sensitive to degradation by prolonged neurotoxic oxidative stress and related inflammation of glial cells (a potential driver of MT), which are responsible for the processing and regulation of cortical iron. Increased iron and decreased MT may thus indicate an adaptive stress response, leading to myelin breakdown (Bartzokis, 2011), such that environmental stress has resulted in elevated neuromelanin levels (Nakane et al., 2008; Chen et al., 2014) to reduce oxidative damage to the hippocampus.

Alternatively, this pattern may be explained by nutritional differences. Glial inflammation is closely linked to iron homeostasis by a complex feedback mechanism, with nutritional iron deficits during foetal, infant, or adolescent development linked to impaired memory and learning (Georgieff, 2008; Carlson et al., 2009, 2010), glial inflammation (Bilbo et al., 2011), and increased risk of neurodegenerative diseases later in life (Gerlach et al., 1994). Iron deficiency exerts numerous effects in the hippocampus and, elevating various metabolites, GABA, and altering overall tissue architecture and myelination (Jorgenson et al., 2005; Georgieff, 2008). Our results might thus indicate an overall neuro-protective effect, in which reduced hippocampal MT (and increased iron) reflect reduced inflammation of neurobiological support structures, caused by either lower trait-stress response or as the result of increased nutritional iron.

Of course, the causal direction and precise mechanisms underlying these effects remain unclear; it could be for example that individuals with a less stressful environment have a better overall memory (and therefore more evidence of neuroprotective element) (Rodrigue et al., 2013), or the iron profiles indicated here may represent a direct response to stress and/or reflect a neuro-genetic ability to adaptively respond to stress or nutritional challenges. We might speculate that metacognition-related hippocampal histology could potentially be an early biomarker for Alzheimer’s and other neurodegenerative diseases; if such a pattern does reflect an early-life response to stress, it may come at the cost of an increased risk to developing these diseases later in life. Regardless, our study provides an interesting starting point for future work, so that we can better understand how the histological factors discovered here relate to metacognitive ability.

### Limitations and Future Directions

Methodological limitations common to all studies requiring spatial normalisation are the potential for residual registration errors, as well as partial volume effects. To minimise these sources of bias we used the DARTEL algorithm for inter-subject registration, which results in maximally accurate registration (Klein et al., 2009), and used the voxel based quantification normalisation procedure to minimise partial volume effects introduced by smoothing (Draganski et al., 2011).

Here we speculatively interpret our results as suggesting a link between the histology of the hippocampus and APFC, and the relationship of memory and metacognition. Indeed, the hippocampus and precuneus are a core part of a memory-related network (Squire, 1992; Brown and Aggleton, 2001). We also found that metacognitive ability both negatively mediated (supressed) the relationship of auditory memory ability and hippocampal/precuneual microstructure, and positively mediated the link between memory and APFC myelination. This suggests that a dynamic interaction between memory and meta-cognition related brain areas mediates the influence of metacognition on memory (or vice versa); however future work with a more general battery of memory and meta-memory tasks is needed to better understand this relationship. Additionally, without stress or nutrition-related measures we can only speculate as to their link with the effects observed here. Indeed, individual differences in brain structure and function are influenced by a variety of developmental, neurogenetic, and environmental factors (Kanai and Rees, 2011).

Thus, an important step for future studies will be to study the microstructural correlates of metacognitive ability with a more comprehensive battery of perceptual, memory, and stress-related measures. Nevertheless, our study is the first to establish that quantitative neuroimaging reveals potential biomarkers sensitive to metacognitive ability, which has allowed us to greatly extend previous functional and volumetric studies. We therefore anticipate that future research will expand upon these findings to better understand their developmental, genetic, and environmental causes.

## Conclusion

By using a quantitative multi-parameter mapping approach, we were able to reveal the contribution of the hippocampus and precuneus to perceptual metacognition. Furthermore, as our method yields standardised quantitative metrics sensitive to the underlying tissue microstructure, these results can be used to inform future clinical research as they can be compared directly across research sites. Our results suggest that the concentration of iron in the hippocampus is an important predictor for metacognitive ability, pointing towards a potential neuroprotective mechanism defending introspection from stress. More generally, these results suggest that memory-related systems may be more important than previously realized for the computation of perceptual confidence. Future research into the genetic, environmental, and other dynamic factors underlying these findings are likely to yield strong dividends in the study of metacognitive ability.

## Acknowledgements

This work was supported by a Wellcome Trust grant 100227 (MA, GR) and an ERC Starting Grant to DSS (310829). The Wellcome Trust Centre for Neuroimaging is supported by core funding from the Wellcome Trust 091593/Z/10/Z.

